# A normalization method that controls for total RNA abundance affects the identification of differentially expressed genes, revealing bias toward morning-expressed responses

**DOI:** 10.1101/2023.10.28.564442

**Authors:** Kanjana Laosuntisuk, Amaranatha Vennapusa, Impa M. Somayanda, Adam R. Leman, SV Krishna Jagadish, Colleen J. Doherty

## Abstract

RNA-Sequencing is widely used to investigate changes in gene expression at the transcription level in plants. Most plant RNA-Seq analysis pipelines base the normalization approaches on the assumption that total transcript levels do not vary between samples. However, this assumption has not been demonstrated. In fact, many common experimental treatments and genetic alterations affect transcription efficiency or RNA stability, resulting in unequal transcript abundance. The addition of synthetic RNA controls is a simple correction that controls for variation in total mRNA levels. However, adding spike-ins appropriately is challenging with complex plant tissue, and carefully considering how they are added is essential to their successful use. We demonstrate that adding external RNA spike-ins as a normalization control produces differences in RNA-Seq analysis compared to traditional normalization methods, even between two times of day in untreated plants. We illustrate the use of RNA spike-ins with 3’ RNA-Seq and present a normalization pipeline that accounts for differences in total transcriptional levels. We evaluate the effect of normalization methods on identifying differentially expressed genes in the context of identifying the effect of the time of day on gene expression and response to chilling stress in sorghum.

## Introduction

RNA-Sequencing (RNA-Seq) is a high-throughput technology for genome-wide transcriptional analysis. RNA-Seq has been widely used in various research areas, including plant biology, to examine many aspects of RNA biology, including differentially expressed genes (DEGs), transcriptome assembly, alternative splicing, variant discovery, cis-regulatory elements, and roles of non-coding RNAs (Stark, Grzelak, and Hadfield 2019). As RNA-Seq was first introduced a decade ago, many advanced RNA-Seq methods have been developed from the standard protocols to answer in-depth issues in molecular biology, especially for determining gene expression at the transcript level. Short-read sequencing is widely used to evaluate DEGs in response to experimental conditions. To prepare samples for short-read sequencing, total RNA is extracted from tissues collected from living organisms. Total RNA comprises 80-90% of ribosomal RNA (rRNAs), while messenger RNA (mRNAs) that include most protein-coding genes make up only 3% (O’Neil, Glowatz, and Schlumpberger 2013). It is necessary to enrich mRNA to increase sequencing efficiency. As RNA-Seq has a complicated workflow, it is easy to introduce biases unintentionally. There are two types of variation: within-sample variation and between-sample variation. GC content, gene length, and contamination are sources of within-sample variation that affect the detection of different genes in the same sample (Evans, Hardin, and Stoebel 2018). Sample preparation and analysis strategies have been developed to target this variation. For example, 3’ RNA-seq reduces the gene length bias by replacing the mRNA isolation and fragmentation steps with cDNA synthesis using oligo-dT primers (Moll et al. 2014).

Identifying biologically relevant between-sample variation is the goal of most RNA-Seq experiments. However, sample preparation techniques can also cause variation between samples. The total number of reads, which reflects sequencing depth, is a critical factor that affects the comparison of gene expression between samples (Evans, Hardin, and Stoebel 2018). Therefore, it is essential to normalize for read depth, so several normalization approaches have been developed to account for biases in the datasets. The built-in normalization method in the two popular differential expression (DE) tools, DESeq2 and EdgeR, normalize gene expression using the distribution of read counts to account for sequencing depth and RNA abundance (Risso et al. 2014; Robinson and Oshlack 2010). Critically, these methods depend on the assumption that most genes do not change in expression between the samples. This fundamental assumption only holds true in some cases. For example, transcriptional amplification in tumor cells globally increases the expression of existing genes rather than turning on the expression of new genes (Lin et al. 2012). Several studies show that experimental conditions alter overall transcription in the cells. In yeast, growth rates are highly correlated with total transcript abundance (Athanasiadou et al. 2016; Brauer et al. 2008; R. Yu et al. 2021), and nutrient limitation also causes transcriptional reprogramming (R. Yu et al. 2020; Lippman and Broach 2009). These cases violate the assumption that the expression level of most genes does not change between samples analyzed by RNA-Seq. Therefore, using the distribution-based normalization in these cases would inappropriately shift the expression levels of genes to force similar total read counts and thus would result in incorrect identification of DEGs (Lovén et al. 2012; Athanasiadou et al. 2016). This means that the total RNA levels must be similar between samples for these distribution-based normalization methods to be appropriate. However, many experimental conditions can change the transcription level or RNA stability, affecting the total RNA abundance, thus invalidating the assumption of consistent total RNA levels.

A related challenge in using distribution-based normalization protocols for RNA-Seq data can happen if genes with a drastic increase or decrease in expression level alter the proportional shares of mRNA in the RNA pool. For example, photosynthesis-related genes, which are about 3000 genes in plants (P. Wang, Hendron, and Kelly 2017), are only up-regulated during the day to perform photosynthesis, and they decrease in expression at night. Assuming that the total RNA pool remains consistent (as most genes are not DEGs), these genes will become a majority in the RNA pool during the day, and normalizing to the same pool size will artificially reduce the abundance of other genes, even those that do not change in expression, leading to incorrect identification of DEGs. Some DEG-identifying algorithms, such as EdgeR, address this for some proportion of the genes (Robinson and Oshlack 2010), but with plants, significant transcriptional changes are frequently induced where photosynthetic genes are a major part of the transcriptome and vary significantly even between different times of day.

Fortunately, there is a simple solution to control for changes in mRNA levels between samples and allow for accurate comparisons. Artificial RNA spike-ins, first developed to account for technical variation in microarray data, can be used as an external control to normalize the total RNA levels (Jiang et al. 2011). These external RNA spike-ins have been proposed since the first high-throughput transcriptional analysis approaches, microarrays, and qRT-PCR (Girke et al. 2000; Hilson et al. 2004; Czechowski et al. 2005). They are often used in RNA-seq studies in mammalian and yeast systems (Brauer et al. 2008; Lun et al. 2017; Byrne et al. 2017; Wilson et al. 2019; Kroustallaki et al. 2019; X. Wang et al. 2021); however, the use of external RNA spike-ins as a normalization factor is not common in plant RNA-Seq analysis studies.

Here, we demonstrate that spike-in normalization is important in identifying DEGs with RNA-Seq. With a synthetic RNA-Seq dataset, we demonstrate that using spike-ins in normalization resulted in better accuracy, specificity, and sensitivity in DEG calling. We demonstrated that adding external RNA spike-ins significantly affects DEG identification between sorghum samples that differ only by the time of day they are collected under normal and chilling stress conditions. We also demonstrate that external RNA spike-ins affect DEG identification between control and chilling stress. Traditional normalization tended to bias toward identifying morning up-regulated genes; however, spike-in normalization reveals novel evening up-regulated genes in sorghum under normal and chilling stress conditions. Our study highlights the importance of spike-in normalization in maintaining wanted variation caused by the experimental conditions and accurately identifying differentially expressed genes.

## Result and Discussion

### 1. External RNA spike-ins can correct for changes in global transcriptional abundance and can capture changes in the composition of differentially expressed genes

Previous studies have demonstrated that RNA spike-ins can accurately correct for changes in global transcription in plants using cDNA arrays, microarrays, and qRT-PCR (Girke et al. 2000; Hilson et al. 2004; Czechowski et al. 2005). Using a synthetic dataset (Figure 1), we demonstrate the effects of a global change in transcription levels of all genes (Figure 1A) or a portion of genes (Figure 1B) on normalization and DEG calling.

**Figure 1.**
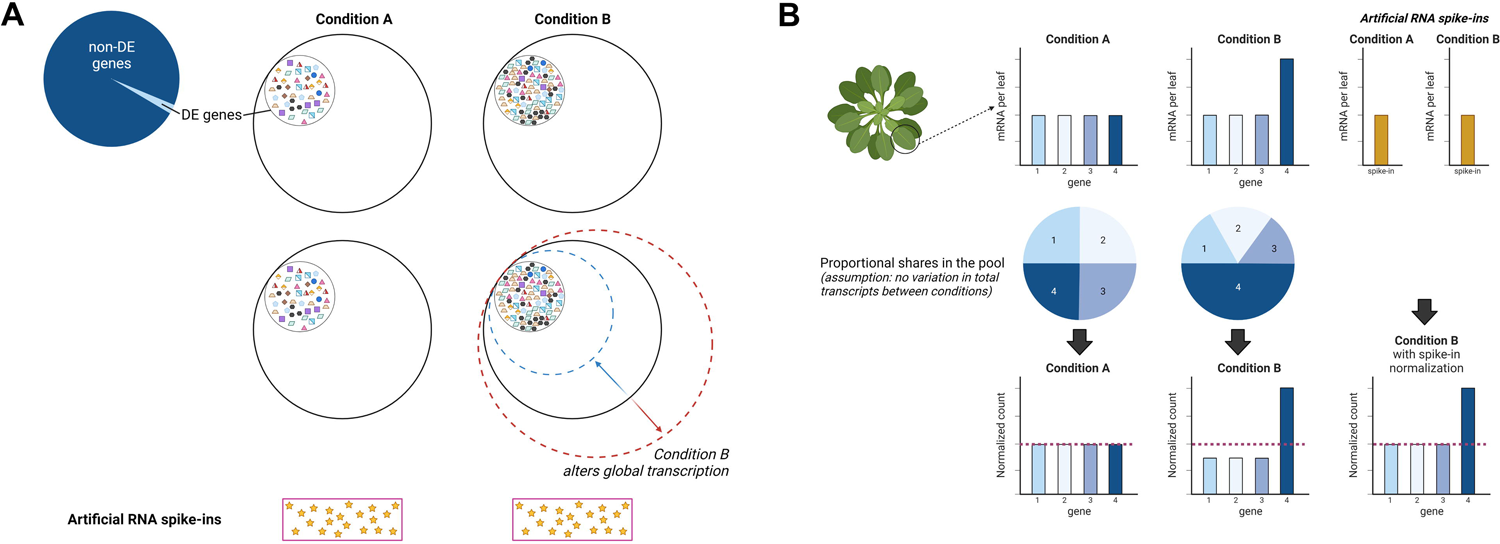
Challenges in RNA-Seq analysis. (A) Commonly-used normalization methods assume that only a small proportion of transcripts are differentially expressed between conditions (small, dashed inner circle in Conditions A and B). It is assumed that most transcripts do not change in expression across experimental conditions (as indicated by solid outer circle in Conditions A and B), resulting in stable transcript pool. However, some experiments do affect global transcription, either increasing (dashed red circle) or decreasing (dashed blue circle) the size of the RNA pool. Figure was created with Biorender.com. (B) A change in proportional share of mRNA in the pool could also affect the identification of DEGs. When some genes substantially increase in gene expression (e.g., Gene group 4), commonly-used normalization methods that assume no change in global expression would result in artificially reducing the expression of other genes (e.g., groups 1-3) that do not change in expression. Figure was created with Biorender.com.

To test the effects of a global change in expression on DEG identification, we created a synthetic RNA-Seq dataset from two experimental conditions, A and B (Figure 2). Each condition had four replicates with varying library sizes, as would be expected from standard library preparation techniques (Figure 2A). In condition B, global transcription is doubled, leading to a doubling of the total reads (Figure 2A). The traditional method employed in DESeq2, known as “Median of Ratio”, calculates the median of the read count ratio in one sample to a geometric mean across all samples (also referred to as a pseudo-reference) (Anders and Huber 2010). To analyze the effects of normalization, we generated relative log expression (RLE) plots, representing the distribution of read counts after normalization (Gandolfo and Speed 2018). Traditional normalization scaled transcript abundance between two conditions to the same level as expected. This indicates that the increased global transcription in condition B is considered noise that needs to be removed (Figure 2A). We tested spike-in normalization using the method Athanasiadou et al. developed (Athanasiadou et al. 2019). In brief, this method utilized a maximum likelihood estimation to determine the calibration constant (□, nu) based on the spike-in read counts (Athanasiadou et al. 2019). Additionally, they computed the library-specific scale factor (□, delta) to account for potential errors introduced during library preparation, assuming that global expression levels between replicates should ideally remain identical (Athanasiadou et al. 2019). The RNA spike-ins were added proportionally to the total sample mRNA according to the experimental protocols for adding spike-ins (Jiang et al. 2011) (Supplemental Table 1). The RLE plot showed increased transcript abundance in the samples in condition B after spike-in normalization (Figure 2A), suggesting that spike-in normalization can detect a change in global transcript abundance. Even when we control the library size to be consistent across samples, we still observe that only spike-in normalization captured a shift in transcript abundance due to altered global transcription (Supplemental Figure 1A). While traditional normalization could not distinguish technical variations introduced by sample preparation from biological differences, the spike-in normalization removed only technical variation but preserved a difference in transcript abundance due to the global change in transcription.

**Figure 2.**
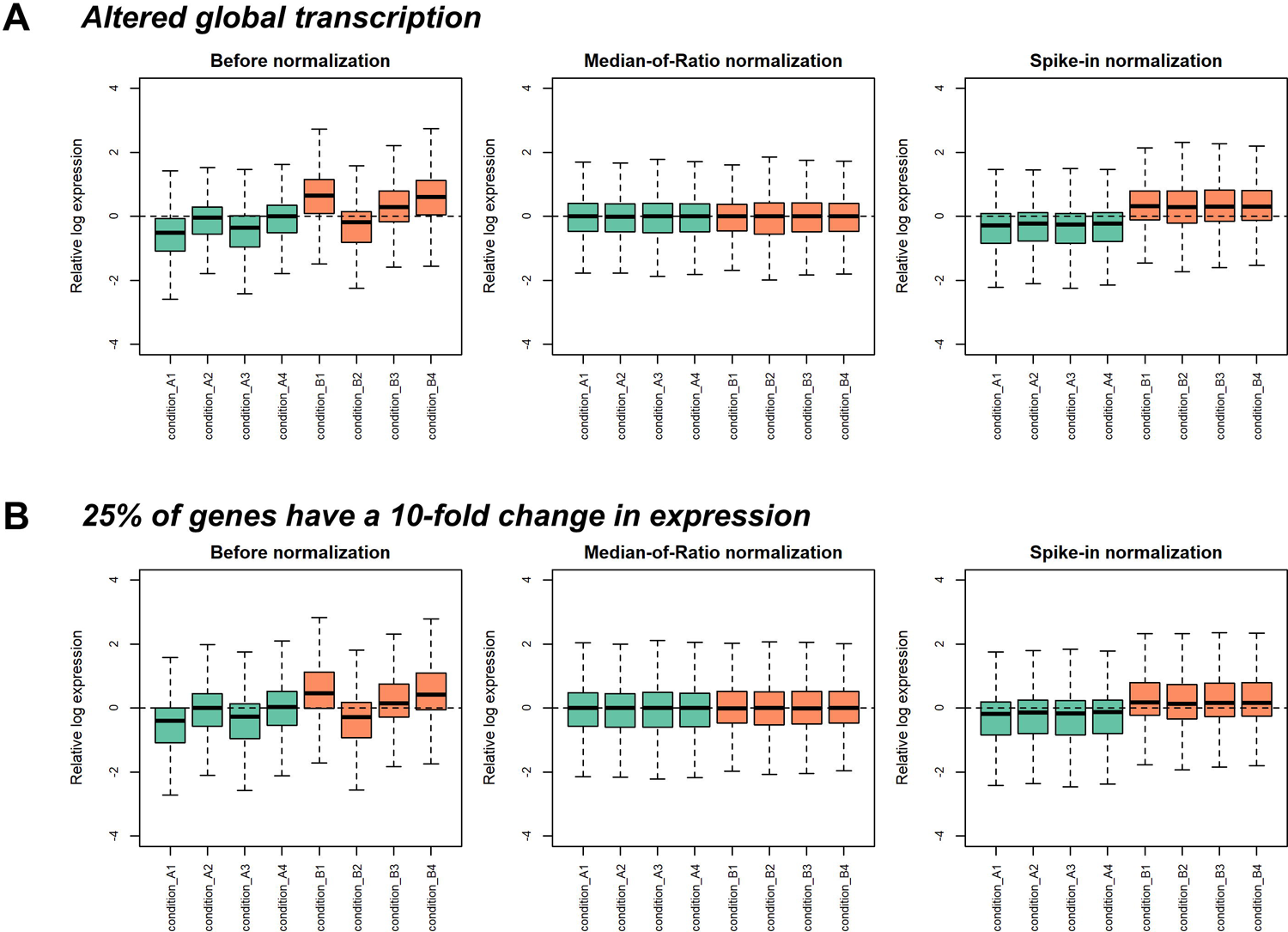
External RNA spike-ins captured difference in transcript abundance due to altered global transcription and altered proportional shares of transcripts. RLE plots of synthetic gene read counts before and after Median-of-Ratio and spike-in normalizations in DESeq2 under altered global transcription (A) and with a drastic change in gene expression in condition B (B) with varied library size across samples.

To assess the impact of spike-in normalization on DEG identification, we created an RNA-Seq dataset comprising 10,000 genes with random numbers generated based on a negative binomial distribution (Supplemental Table 2). Under conditions with a substantial increase in gene expression, the traditional normalization adjusted the read count distribution to be uniform across the samples (Figure 2B and Supplemental Figure 1B to 1D). In contrast, the spike-in normalization method effectively maintained the unequal distribution between conditions A and B (Figure 2B and Supplemental Figure 1B to 1D). For example, in scenarios where 25% and 50% of genes were up-regulated in condition B, more DEGs identified by the spike-in normalization method overlapped with the list of genes we manipulated to be DEGs than those identified by the traditional method (Supplemental Table 3).

To further assess the performance of the normalization methods, we constructed a confusion matrix, which allowed us to compare the predicted DEGs (after normalization) with the expected DEGs (genes for which we manually adjusted the expression). The spike-in normalization method exhibited more true positives and true negatives, leading to higher accuracy, sensitivity, and specificity (Supplemental Table 3), suggesting that external RNA spike-ins significantly enhance DEG identification when drastic changes in gene expression affect the proportional shares of mRNA in the pool. This demonstrates the effectiveness and robustness of the spike-in normalization method in improving the accuracy and reliability of DEG analysis in diverse experimental conditions.

### 2. Using external RNA spike-ins leads to identifying novel PM up-regulated genes in sorghum

Through evaluation of a synthetic RNA-Seq dataset, we observed that spike-in normalization significantly enhances the accuracy of DEG identification. However, to validate the performance of spike-in normalization in a real RNA-Seq dataset, we explored its effectiveness in understanding the impact of time of day and chilling stress on gene expression in sorghum leaves. We utilized 3’ RNA-Seq to analyze the transcriptomic changes in sorghum leaves under control and chilling stress conditions during both morning and evening time points. The utility of 3’ RNA-Seq in studying transcriptional responses has been demonstrated in various plant species, including maize, rice, *Brachypodium distachyon*, *Setaria viridis*, switchgrass, apples, tomatoes, and Alpine orchid (*Gymnadenia rhellicani*) (Eveland, McCarty, and Koch 2008; Kremling et al. 2018; Israeli et al. 2019; Palmer et al. 2019; Silva et al. 2019; Kellenberger et al. 2019; Y. Yu et al. 2020). One of the advantages of 3’ RNA-Seq is its ability to minimize gene length bias by capturing only one read per transcript (Ma et al. 2019). The success of 3’ RNA-Seq relies heavily on the quality of the reference genome (Tandonnet and Torres 2017; Ma et al. 2019). Encouragingly, we found that 88% to 92% of the reads uniquely mapped to the sorghum BTx623 reference genome (Supplemental Table 4). The reads predominantly aligned towards the 3’ end of the gene body, which is consistent with our expectations and the characteristics of 3’ RNA-Seq (Supplemental Figure 2A), suggesting that the sorghum genome quality is sufficient for 3’ RNA-Seq applications.

Artificial RNA spikes (SIRV set 3, Lexogen, USA) were incorporated into the plant samples during the RNA extraction process. The SIRV set 3, comprising ERCC spike-in controls <u>(Lemire et al. 2011)</u> and Lexogen’s Spike-In RNA variants (SIRVs) controls <u>(Paul et al.</u> <u>2016)</u>, is a well-established tool commonly used in mammalian and yeast studies (Blevins, Carey, and Albà 2019; Nadal-Ribelles et al. 2019; Topal et al. 2019) to account for variations in total RNA content between samples. While in mammalian cells, spike-in standards are added proportionally to the cell count (Lovén et al. 2012), determining the ideal method for adding RNA spike-ins to plant samples presents unique challenges. Unlike mammalian cells, accurately assessing the total cell number in plant tissue is more complicated. Nonetheless, several approaches have been considered, such as normalizing to the plant, leaf, tissue weight, or total DNA content. In chilling stress, our focus, changes in cell expansion, cell cycle progression, and potential endoreduplication issues (Rymen et al. 2007; Louarn, Andrieu, and Giauffret 2010; Zhao et al. 2014; Ashraf and Rahman 2019) have been observed. Therefore, we opted to normalize on a per-leaf basis, adding 90 pg of spike-in controls to each sorghum leaf sample, where each sample originated from a single leaf. While chilling stress resulted in a noticeable reduction in leaf size, no size difference was observed between morning (AM) and evening (PM) samples collected on the same day. Therefore, to initially assess the efficacy of spike-ins, we focused on identifying DEGs between the AM and PM samples in either control or chilling conditions so that the leaf size would not be a factor (Figure 3). Notably, the number of spike-in reads correlated with total read counts (Supplemental Figure 2B), and the proportion of spike-in reads to total reads remained consistent across samples, representing approximately 0.04 - 0.08% (Supplemental Figure 2C). A one-way ANOVA was performed to evaluate treatments’ impact on the spike-in proportion, resulting in a p-value (0.336) higher than 0.05 (Supplemental Figure 2C), indicating no significant difference in the spike-in proportion between treatments, as expected.

**Figure 3.**
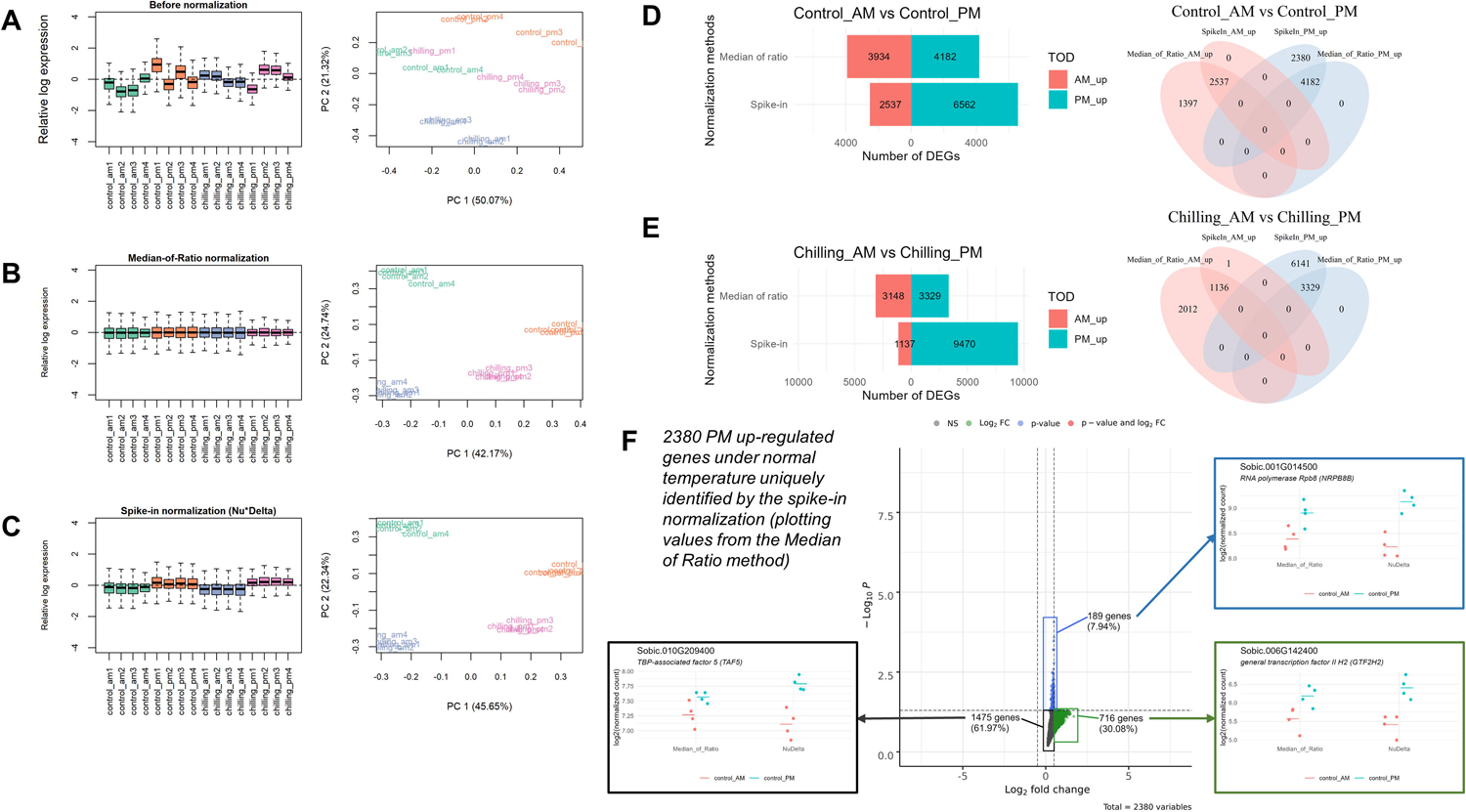
External RNA spike-in normalization preserves variation between experimental conditions. (A to C) RLE and PCA plots of unnormalized read counts (A), read counts after the Median-of-Ratio normalization (B), and read counts after the spike-in normalization method in DESeq2 (C). (D to E) Bar charts and Venn diagrams show the number of morning-upregulated (Salmon) and evening-upregulated (Teal) genes undernormal (D) and chilling stress (E) from the Median of Ratio and spike-in normalization methods. Genes with FDR < 0.05 and the absolute log_2_ fold-change > 0.5 were identified as differentially expressed genes (DEGs). (F) Volcano plot showing the 2380 genes identified as DEGs by spike-in method under the control condition. These genes are plotted with their values after the Median of Ratio normalization to visualize why they were not identified as DEGs by the Median of Ratio method. Dot plots represent an example gene for each category not identified as DEGs by the Median of Ratio due to failed log_2_ fold-change (blue), failed p-value (green), and failed log_2_ fold-change and p-value (black).

We employed RLE plots to visualize the transcript distribution in our data (Figure 3). As expected, before normalization, the boxes on the plots exhibited noticeable deviations from the center (Figure 3A). The PCA plot illustrated that the biological replicates of each treatment were not well clustered (Figure 3A), indicating the need for normalization before proceeding with the differential gene expression analysis. The traditional normalization approaches completely removed the variation between biological and technical replicates, aligning all samples to the same level, as anticipated, based on their underlying assumption (Figure 3B). In contrast, reads normalized using the normalization factors derived from the spike-in read count reduced the variation within biological replicates while retaining the variation between treatments (Figure 3C). The PM samples generally displayed a higher average read count than the AM samples under control and chilling stress conditions (Figure 3C). Despite differences in leaf size, we noticed little difference in the average read count when comparing AM control to AM chilling or PM control to PM chilling (Figure 3C). Additionally, including spike-ins reduced the within-treatment variation between samples in the same condition compared to the traditional method (Figure 3C). This suggests that the use of spike-in normalization not only impacted between-treatment total read levels but also reduced variation within samples of the same condition.

The PCA plots for both traditional and spike-in-based normalization demonstrated that the first principal component distinctly separated samples based on the time of day, while the second component effectively distinguished samples according to the temperature condition (Figure 3B and 3C). These findings suggest that the time of day exerts a more significant influence on gene expression variation in leaves of this sorghum genotype than the differences between the control and chilling temperatures. Consequently, this highlights the critical importance of evaluating stress responses multiple times throughout the day to obtain a comprehensive and accurate representation of gene expression changes <u>(Grinevich et al. 2019;</u> <u>Fowler and Thomashow 2002; Blair et al. 2019; Fowler et al. 2005; Bonnot et al. 2023)</u>.

We conducted DEG analysis between dawn and dusk in both the control (control_AM vs control_PM) and chilling temperature (chilling_AM vs chilling_PM) conditions to assess the effect of time of day on gene expression in sorghum under two temperature conditions. Employing the RNA spike-ins for normalization identified a significantly higher number of up-regulated genes in the evening (Figure 3D and 3E). This increase in the identification of evening up-regulated genes by normalization methods that included spike-ins compared to traditional normalization methods was robust across several spike-in and traditional normalization methods (Supplemental Figure 3A and 3B). To compare the effects of adding RNA spike-ins, we focused on comparing the DESeq normalization without using RNA spike-ins (Median of Ratios) (Anders and Huber 2010) to the RNA spike-in adjusted (Athanasiadou et al. 2019). The DEGs exclusively detected by the Median of Ratio method were all morning up-regulated (Figure 3D and 3E). Conversely, the spike-in normalization uniquely identified novel evening up-regulated genes in both temperature conditions (Figure 3D and 3E). This observation suggests that the traditional normalization approach, which adjusts the transcript abundance distribution to the same level, introduces a bias toward detecting morning up-regulated genes (Figure 3D and 3E). In contrast, normalizing the data with external RNA spike-ins successfully maintained the asymmetric transcript distribution in total reads between morning and evening samples (Figure 3D and 3E). This preservation of asymmetric distribution facilitated the detection of more evening up-regulated genes (Figure 3C), further emphasizing the advantages of utilizing RNA spike-ins for accurate normalization and more comprehensive gene expression analysis in our study.

### 3. Genes uniquely identified by the addition of RNA spike-ins are due to differences in both assigned log fold changes and significance

To investigate the reasons behind the differences in the DEGs identified by the Median of Ratio and RNA spike-in normalization methods, we visually examined the log_2_ fold-change and p-values of the AM vs. PM DEGs exclusive to each method (Figure 3F and Supplemental Figure 3C to 3E). Under the control conditions, we found that 1397 unique DEGs from the Median of Ratio method were up-regulated in the morning, while all 2380 unique DEGs from the spike-in method were up-regulated in the evening (Figure 3D). Examining why the DEGs identified by the RNA spike-in normalization method were not identified in the Median of Ratio method showed that of these 2,380 genes, a few were not identified by the Median of Ratio method because they did not make the fold-change cutoff (7.94%) or the p-value cutoff (30.08%). However, most of the DEGs uniquely identified (61.97%) failed to make either cutoff when using the Median of Ratios (Figure 3F). A similar distribution was observed for the DEGs uniquely identified by the Median of Ratio method. Of these 1,397 DEGs, 10.59% passed the p-value threshold but did not meet the fold-change cutoff; 24.7% passed the fold-change cutoff but not the p-value; and the majority of the DEGs identified only by the Median of Ratio method, 64.71% did not meet either the p-value or fold-change cutoff in the RNA-spike in normalization (Supplemental Figure 3C).

We plotted the expression of selected genes in each category to visualize why the normalization method would make a difference in log fold-change, p-value, or both. We observed that both the variation between the biological replicates and the expression level in the PM samples contributed to these differences (Figure 3F and Supplemental Figure 3C). For example, both an increase in average expression in the evening and a reduction in within-condition variation contributed to the genes uniquely identified by the RNA spike-in normalization (e.g., *GENERAL TRANSCRIPTION FACTOR II H2* (*GTF2H2*)*, RNA POLYMERASE RPB8* (*NRPB8*), and *TBP-ASSOCIATED FACTOR 5* (*TAF5*)) (Figure 3F). For some genes (e.g., *NRPB8* and *TAF5*), there also appears to be a reduction in the average expression of the AM timepoint after the RNA spike-in normalization. Examples of genes uniquely identified as DEGs and expressed higher in the AM than in the PM by the Median of Ratio method include *BASIC LEUCINE-ZIPPER 43* (*bZIP43*)*, AUXIN RESPONSE FACTOR 16* (*ARF16*), and *PSEUDO-RESPONSE REGULATOR 7* (*PRR7*) (Supplemental Figure 3C). For these three genes, the RNA spike-in normalization reduced the within-condition variation between samples and increased the expression of each sample in the PM, thus reducing the statistical significance of their difference (*bZIP43*), total change in expression levels (*ARF16)*, or both (*PRR7)*.

The improvement of transcript abundance with the spike-in normalization was also observed in the AM vs. PM DEGs under chilling stress conditions. Over 80% of non-DEGs did not pass both p-value and log fold-change cutoffs (Supplemental Figure 3D and 3E).

Furthermore, some genes in this group showed an opposite expression direction in the chilling stress conditions (Supplemental Figure 3D and 3E). For example, Sobic.003G048600 (*GLUTAREDOXIN 4*, *GRX4*) and Sobic.003G257600 (*TAF11*) were significantly up-regulated in the evening with the spike-in normalization method, but their expression increased in the morning with the default normalization (Supplemental Figure 3D). Sobic.001G537300 (*HOMEOBOX-LEUCIN ZIPPER PROTEIN REVOLUTA*, *REV*) and Sobic.007G186300 (*ANKYRIN REPEAT-CONTAINING PROTEIN 2*, *AKR2*) were significantly up-regulated in the morning with the default normalization (Supplemental Figure 3E). However, they tended to be down-regulated in the morning with the spike-in normalization normalization (Supplemental Figure 3E). In addition, the biological replicates in the evening samples after the RNA spike-in normalization were more tightly grouped with each other than the replicates after the Median of Ratio method (Supplemental Figure 3D and 3E). These findings underscore the importance of carefully selecting and applying appropriate normalization methods to accurately interpret and compare gene expression patterns, as the normalization method affects not only the statistical significance, but can also alter the log fold-changes differences between treatments.

### 4. Functional analysis indicates that normalization with RNA spike-ins changes the enriched cellular functions identified

We demonstrated that the DEGs uniquely identified by the Median of Ratio method exhibited a bias toward morning up-regulated genes. In contrast, the RNA spike-in method revealed more unique evening up-regulated genes. To gain insights into the cellular functions of DEGs detected by both methods and those uniquely identified by each method, we conducted MapMan analysis to categorize the genes (Figure 4). In the control condition, both methods yielded a comparable number of unique DEGs falling into cellular pathways such as cell organization, development, hormones, regulation, and redox (Figure 4A). However, the RNA spike-in method notably showed more significant enrichment of the identified DEGS than the Median of Ratio method in the following pathways: DNA repair, cell cycle, protein targeting, biotic stress response, RNA synthesis, RNA processing, and protein synthesis (Figure 4A). In the context of DEGs between AM and PM under chilling stress conditions, the RNA spike-in method exhibited significant enrichment in almost all cellular pathways compared to the DEGs uniquely detected by the Median of Ratio method (Figure 4B). These results indicate a common function of the DEGs uniquely identified in the RNA spike-in method. Since the RNA spike-in identified genes are all higher expressed in the PM samples, these results support the idea that evening-specific functions and cellular processes may be under-represented or missed using traditional DEG analysis.

**Figure 4.**
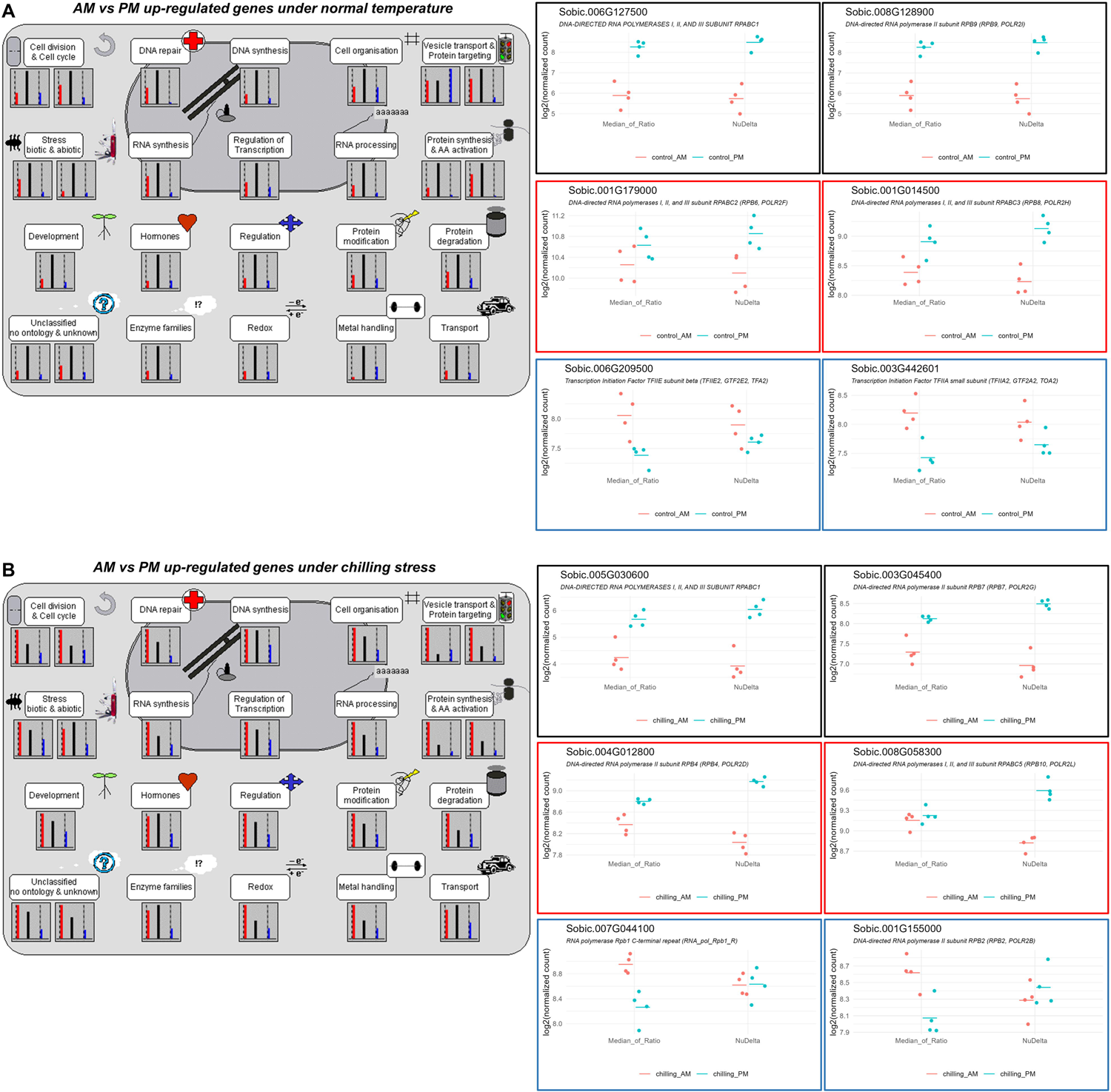
Spike-in normalization-specific DEGs were enriched in various biological pathways including RNA synthesis. MapMan classified DEGs between morning and evening in the control (A) and chilling stress conditions (B) identified by the Median of Ratio method only (blue, right bar), by spike-in only (red, left bar), and by both methods (black, center bar) into different cell functional groups. Dot plots represent examples of genes from the RNA synthesis group after different normalization methods.

We focus on genes in RNA synthesis, a fundamental process in the central dogma of gene expression. In MapMan analysis, we found that the RNA spike-in method identified more unique DEGs relating to RNA and protein synthesis than the Median of Ratio method under control and chilling stress conditions (Figure 4). In the control condition, 534 RNA-related DEGs were identified by both normalization methods, and 128 and 227 genes were unique to the Median of Ratio and RNA spike-in methods, respectively. In the chilling stress condition, there were 402 common DEGs in both methods, and 215 and 530 genes were uniquely found in the Median of Ratio and RNA spike-in methods, respectively. RNA polymerases are essential for RNA production in cells, especially RNA polymerase II which is responsible for mRNA synthesis <u>(Kwapisz et al. 2008; Vannini and Cramer 2012)</u>. Both the Median of Ratio and RNA spike-in methods identified that several nuclear RNA polymerase (*NRP*) genes were up-regulated at night under the control condition, and spike-in normalization method identified additional evening up-regulated *NRP* genes which were not detected as DEGs with the traditional normalization (Figure 4 and Supplemental Table 5). Even more *NRP* genes were up-regulated in the evening under chilling stress with the spike-in normalization. Interestingly, Sobic.007G044100 (*NRPB1*) and Sobic.001G155000 (*NRPB2*), the largest catalytic subunits in the RNAP II that are commonly shared among RNAP II, IV, and V and function in the catalytic site <u>(Ream et al. 2014)</u> were identified as morning up-regulated genes under chilling stress by the traditional normalization. In contrast, these two genes were not differentially expressed by the spike-in normalization (Figure 4 and Supplemental Table 5). Although only measured at the transcriptional level, this increase in the abundance of RNAP II subunits might indicate that RNAP II is more active, or, if these are subunits shared with RNAP IV, that RNAP IV is more active, supporting why there is an observed increase in global transcripts when using the RNA spike-in method. Several publications indicate that the expression of genes encoding the RNAP subunits is affected by abiotic stress and some are required for abiotic stress tolerance <u>(Fernández-Parras et al. 2021; Popova et al. 2013; Borsani et al. 2005)</u>. This provided a possibility that the sorghum produces more RNAP subunits at night to reprogram gene expression in response to temperature changes, and the RNA spike-in normalization identified additional nighttime up-regulated RNAP coding genes.

### 5. Normalization with RNA spike-ins affects the evaluation of the response to chilling stress

Based on our evaluation of the differences between the normalization methods, we evaluated the response to chilling stress compared to the control conditions in sorghum at dawn and dusk using both normalization methods. DE analysis showed that the traditional normalization method detected a comparable number of chilling up- and down-regulated genes in the morning (3411 and 3739 genes, respectively) and evening (2309 and 2611 genes, respectively) (Figure 5A and 5C). In contrast, RNA spike-in normalization identified more chilling down-regulated genes (4571 down-regulated and 2877 up-regulated genes) in the morning but more chilling up-regulated genes in the evening (3478 up-regulated and 1607 down-regulated genes) (Figure 5A and 5C). Other spike-in utilized methods exhibited similar results where more chilling down-regulated genes were detected in the morning and chilling up-regulated genes (Supplemental Figure 4A and 4B). The biggest factor causing genes to be classified in opposite expression is that the spike-in method shifted the mean of expression downward for the AM chilling samples and upward for the PM chilling samples (Figure 2C). Heatmaps of 1160 genes down-regulated by chilling in the morning that were identified as DEGs by the spike-in normalization clearly showed a difference in the chilling samples. The expression of these genes in chilling conditions after the RNA spike-in normalization was lower than the expression after the default normalization (Figure 5B). Likewise the difference was distinctly in the chilling samples for the 863 AM chilling up-regulated DEGs uniquely identified the Median of Ratio. These genes are identified as more highly expressed under chilling samples by the Median of Ratio method than by the RNA spike-in method. These results suggests that there is a bias in the Median of Ratio normalization for identifying up-regulated genes in the morning.

**Figure 5.**
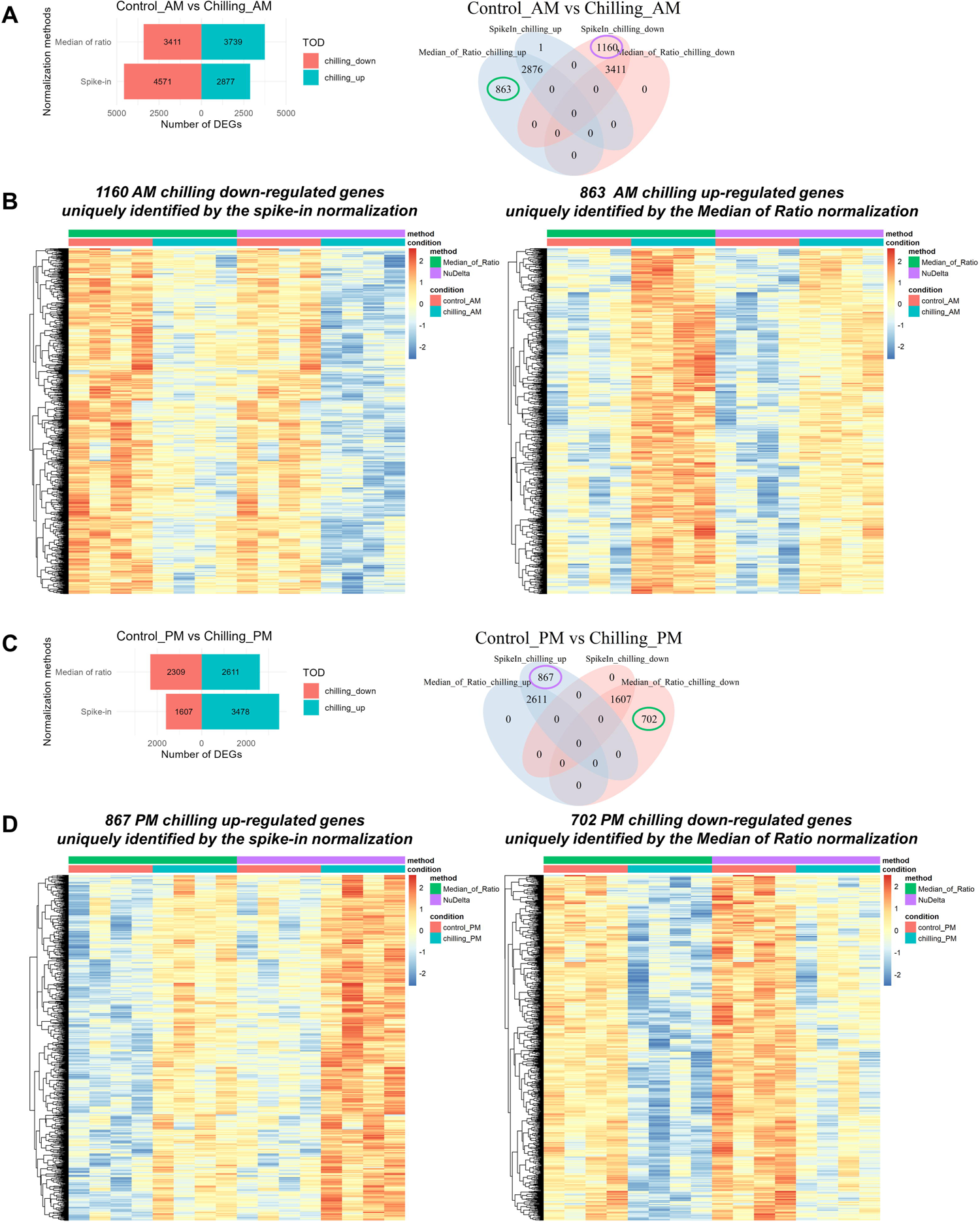
Spike-in normalization revealed contrasting responses to chilling stress at different times of the day. Spike-in normalization identified more genes as down-regulated chilling DEGs genes in the morning but more up-regulated genes in response to chilling stress in the evening. Bar charts and Venn diagrams (A and C) show the number of chilling down-regulated and up-regulated genes in the morning (A) and in the evening (C) from the Median of Ratio and spike-in normalization methods. Genes with FDR < 0.05 and the absolute log fold-change > 0.5 were identified as differentially expressed genes. (B and D) Heat maps display normalized expression of chilling down-regulated and up-regulated genes in the morning (A) and in the evening (C) that were uniquely identified by the spike-in (left, green top bar) and Median of Ratio (right, purple top bar) methods.

Furthermore, in the evening (PM) samples, the 867 genes up-regulated under chilling that were uniquely identified by the RNA spike-in method had a strong increase in expression under chilling compared to the expression normalized by the traditional, Median of Ratio method (Figure 5C). In contrast, the expression of the 702 PM DEGs in the evening identified by the Median of Ratio method as down-regulated under chilling method were both higher under control conditions when normalized by the RNA spike-in method and lower under chilling conditions in the Median of Ratio method analysis (Figure 5D). Thus, in the evening samples, the Median of Ratio method is identifying fewer chilling up-regulated genes and more down-regulated genes.

Gene Ontology (GO) enrichment analysis of chilling up- and down-regulated genes in the morning showed that both normalization methods shared several similar GO terms, for example, cell redox homeostasis, protein folding, response to endoplasmic reticulum stress showed up in chilling down-regulated genes, and metabolic process, response to cold, chloroplast organization, and carbohydrate metabolic process showed up in chilling up-regulated genes (Supplemental Figure 5C and 5D). In the evening, chilling down-regulated genes were enriched in GO terms related to translation while chilling up-regulated genes were enriched in photosynthesis and light response (Supplemental Figure 5E and 5F). However, there are numerous GO terms that appeared in one normalization method but not in another (Supplemental Figure 5C to 5F). For example, photorespiration is identified as enriched in genes down-regulated by chilling in the morning when normalized with RNA spike-ins but not the Median of Ratio method (Supplemental Figure 5C). Our results indicated that the differences in DEGs captured by the RNA spike-in normalization, propigates into unique GO terms in response to chilling stress and could affect our understanding in chilling stress response in plants.

### 6. The implication of RNA spike-in controls in RNA-Seq normalization

We have demonstrated that spike-in normalization significantly impacts differential expression analysis, which is a crucial aspect of RNA-Seq studies. Based on our synthetic data, spike-in normalization provides more accurate results, especially in cases where global transcription and proportional shares of the RNA pool are affected by experimental conditions. Several environmental changes have been shown to have effects on global transcription in plants (Branco-Price et al. 2005; Szádeczky-Kardoss et al. 2022; Rymen et al. 2007; T. Zhang et al. 2012; H. Zhang et al. 2022; Koiwa 2006; Thatcher et al. 2018). Although artificial spike-in controls have been used in microarray and some RNA-Seq data, their application in plant RNA-Seq data has been relatively limited. Some RNA-Seq protocols incorporate normalization factors such as spike-ins during library preparation to account for variation in library size.

However, we observe that adding spike-ins only as a control for library variation has an overall reduction in the DEGs identified in each contrast (e.g., DESeq2 Median of Ratios vs DESeq2 RUV and EdgeR TMM vs EdgeR RUV in Supplemental Figures 3 and 5) but no effect on the overall proportion of genes identified as DEG in either contrast. Normalization approaches that incorporate RNA spike-ins added during RNA extraction to account for changes in total transcript levels have an effect on the distribution of DEGs identified (Supplemental Figures 3 and 5). Thus, considering the potential effects of experimental conditions on transcription in organisms, it is essential to consider using spike-ins in RNA-Seq experiments to ensure reliable and interpretable results.

In our sorghum dataset, spike-in normalization revealed that more transcripts were accumulated at night than at dawn in this genotype, irrespective of temperature conditions, suggesting that the regulation of transcription machinery by the circadian clock possibly occurs in plants, similar to mammalian studies (Koike et al. 2012; Y. Wang et al. 2018; Doherty and Kay 2012). For example, studies in mice using ChIP-Seq of RNA pol II in preinitiation and elongation states have indicated that RNA pol II activity depends on the time of day (Koike et al. 2012). The nascent transcription peaks at night (CT14.5). At the same time, the active form of RNAPII with phosphorylation of Serine 5 at the C terminus domain (CTD) highly accumulates on the genome in the early morning (CT0.6), suggesting that nascent transcription starts at night. Other histone modifications, such as promoter-enriched H3K27ac and elongation marks H3K36me3 and H3K79me2, also peak after the nascent transcription peak at night. Interestingly, the amount of nascent transcripts highly accumulate around CT15, but it is not highly correlated with the accumulation of mature mRNAs, with the mean circadian phase around CT7. This suggests that post-transcriptional modifications also play a role in driving the rhythmicity of mRNA levels.

Various publications have identified genes with rhythmic expression under diurnal conditions in different plant species (Michael et al. 2008). For instance, in Arabidopsis, approximately 89% of transcripts cycle when plants are grown under photocycles, thermocycles, or combined photocycles and thermocycles (Michael et al. 2008). Similarly, sorghum, maize, and foxtail millet show that 52%, 30%, and 43% of genes exhibit rhythmic expression under diurnal conditions (Lai et al. 2020). Furthermore, around 60% and 59% of transcripts show rhythmic expression in poplar (*Populus trichocarpa*) and rice (*Oryza sativa* ssp. japonica) (Filichkin et al. 2011). These findings strongly suggest a circadian regulation of transcription in plants. Additionally, post-transcriptional modifications, such as alternative splicing and polyadenylation, also occur in a time-of-day-dependent manner in plants.

By employing RNA spike-in normalization, we preserved the asymmetric distribution of transcript abundance influenced by the time of day in the differential expression analysis. This analysis using RNA spike-ins resulted in identifying more differentially expressed genes, especially those up-regulated in the evening. It indicates that the current RNA-Seq analysis workflow may not fully capture the variation in total RNA levels when comparing morning and evening samples, potentially leading to an artificial reduction in evening gene expression to align it with the average gene expression in the morning. This, in turn, reduces the ability to identify genes expressed at higher levels at night compared to dawn. However, by utilizing artificial RNA spike-ins and normalization methods that leverage their benefits, we improve the identification of genes expressed during the nighttime. Analyzing both control and chilling stress conditions with the spike-ins reveals that several biological processes, even those previously unknown to have time-of-day-specific regulation, were expressed at higher levels at night. Critically, transcription and translation, which are fundamental components of the central dogma show significant time-of-day specific regulation emphasizing the potential need to control for global changes in gene expression.

## Materials and Methods

### 1. Plant growth conditions and sample collection

*Sorghum bicolor* cultivar BTx623 was grown in a controlled environment chamber (Conviron model PGR15; Winnipeg, MB, Canada) facility at the Department of Agronomy, Kansas State University. The chambers were maintained at 12 h photoperiod, with 800 μmol m^-2^ sec^-1^ light intensity at 5 cm above the canopy and 60% relative humidity (RH). The daytime/nighttime temperatures for the controlled and chilling stress conditions were 30/20°C and 20/10°C, respectively, in the growth chambers with the 12/12 hours of light/dark cycles. The chambers were programmed to reach the daytime (0800 to 1700 h) target temperatures of 30 and 20°C, by following a gradual increase from 20 to 30°C (control) and 10 to 20°C (chilling stress) respectively, with a 3 h transition (0500 to 0800 h). Similarly, the nighttime (2000 to 0500 h) target temperatures of 20 and 10°C was obtained by a gradual decrease in temperature from 30 to 20 °C (control) and 20 to 10°C (chilling stress) respectively, with a 3h transition (0500 to 0800h). The second fully-expanded leaf with a visible collar from the top on the 15-day-old seedlings were collected at 1 hour after the light was on as the morning samples and at 1 hour before the light was off as the evening samples. The fresh samples were flash-frozen in liquid nitrogen and stored at −80°C.

### 2. RNA extraction and library preparation

Total RNA was extracted from leaf tissue using the RNeasy Plant Mini Kit (Qiagen, USA). 5.94 µL of 15.15 pg/µL Spike-in RNA variant controls (SIRVs) set 3 (Lexogen, USA) were added after tissue grinding to obtain 90 pg of SIRV per leaf. DNase I treatment was performed on RNeasy spin columns during a washing step. Total RNA was measured by Nanodrop Lite spectrophotometer (Thermo Scientific, USA). 300 ng of total RNA was used in the QuantSeq 3’mRNA-Seq Library Prep Kit for Illumina (FWD) (Lexogen, USA) according to a manufacturer’s instruction. In brief, oligo (dT) primers were used to isolate mRNA and to start the first-strand cDNA synthesis. After that, RNA was removed, and the second-stranded cDNA synthesis was performed using random primers containing Illumina-compatible linker sequences. cDNA libraries were purified with magnetic beads. Library amplification with 18 rounds of PCR was performed using i7 index primers. Libraries were measured and quality checked by an Agilent 2200 TapeStation. 10 nM pooled libraries were sequenced by Illumina NovaSeq 6000 platform with 100-bp single reads at NC State University Genomic Sciences Laboratory (Raleigh, NC, USA).

### 3. RNA-Sequencing data processing

Raw reads were quality checked by FastQC (version 0.11.8) (Andrews 2010). Adapter trimming was performed by BBDuk (in BBMap version 38.34) with the following parameters: *k=13, ktrim=r, useshortkmers=t, mink=5, qtrim=r, trimq=10, minlength=20* (Bushnell 2014; “BBMap Guide” 2016). FastQC was used again to check the reads after trimming. Reads were mapped to Sorghum bicolor cultivar BTx623 reference genome (Phytozome 12, genome version 3.1.0 and annotation version 3.1.1) (McCormick et al. 2018) and SIRV sequences (Lexogen, USA) using STAR (version 2.5.3) with default parameters (Dobin et al. 2013). SAMtools (version 1.9) was used to sort and index BAM files. Read count tables were generated using HTSeq-count in the HTSeq package (version 0.11.2) (Anders, Pyl, and Huber 2014). Low read counts were filtered out using the *filterByExpr()* command in EdgeR, resulting in 17651 genes for further analysis (Robinson, McCarthy, and Smyth 2010). With the sequencing read depth we used, about 60% of spike-in transcripts were detected with 3’ RNA-Seq, with 58 out of 92 ERCC and 42 out of 68 SIRV transcripts being identified. The non-detected ERCC transcripts mostly had concentrations lower than 2 amoles/ul (Supplemental Figure 5A), suggesting that our sequencing approach would likely miss transcripts with concentrations below this limit. This provides a valuable quantitative limit of detection not commonly available in traditional RNA-Seq studies.

### 4. Normalization prior to differential expression analysis

Normalization was performed before a differential expression analysis in DESeq2 and EdgeR (Supplemental Figure 5B). Traditional normalization in DESeq2 was a Median of Ratio method (Anders and Huber 2010). In DESeq2, we used the command *DESeq()* to perform normalization and differential expression analysis at the same time. EdgeR has a Trimmed Mean of M values (TMM) method as a default normalization <u>(Robinson and Oshlack 2010)</u>. The command *calcNormFactors(method = “TMM”)* was run to perform TMM normalization in EdgeR.

To normalize gene read counts based on spike-in reads in DESeq2, we used the method developed by Athanasiadou et al. (2019), and this method is referred to as “spike-in normalization” throughout this manuscript and “DESeq2 Spike-in (Nu*Delta)” in Supplemental Figures 3, 4, and 5. This method estimates calibration constants based on the abundance of spike-in controls and library-specific correction factors accounting for unwanted variations. The maximum likelihood calibration constant (*v_j_*, Nu) was estimated from three factors: (1) the proportion of spike-in counts across all libraries contributed by the reference spike-in, the molecules per sample for the reference spike-in and the size of spike-in library *j* (Athanasiadou et al. 2019). Then the nominal abundance of transcripts in library *j* is the transcript counts divided by *v_j_* (Athanasiadou et al. 2019). The library-specific scaling factor (δ*j*, Delta) is based on the assumption that the transcript abundance should be identical across technical replicates in each treatment (Athanasiadou et al. 2019). δ*j* is the scaling factor for library *j* in condition *l* is the exponential of β*_j_*where β*_j_*is the difference between the mean of log(nominal abundance of transcript *i* in library *j*) and the mean of log transformation of nominal abundance of transcript *i* among libraries in condition *l* (Athanasiadou et al. 2019). In DESeq2, *v_j_*and δ*j* were used as a size factor given by *v_j_*δ*_j_* divided by a geometric mean of *v*δ (Athanasiadou et al. 2019). We also tested other methods utilizing spike-in read counts in DESeq2 were from Brennecke et al. (2013). The Median of Ratio method was used to calculate size factors from the spike-in read count table by running *estimateSizeFactors()* and *sizeFactors()* commands <u>(Brennecke et al. 2013)</u>. The size factor values were then added to the gene read count dataset prior DE analysis This method is referred to as “DESeq2 Spike-in size factor” in Supplemental Figures 3, 4, and 5.

We utilized a spike-in based normalization in EdgeR. We applied a log_2_ transformation to total spike-in reads to obtain normalization factors of each library and stored them as offsets to normalized gene counts during the DE analysis <u>(Lun et al. 2017)</u>. This is identified as EdgeR log_2_ spike-in in Supplemental Figures 3, 4, and 5. As a contrast, we used the spike-ins to correct for library size only (ignoring changes in global gene expression). We did this using Removed Unwanted Variation (RUV) to calculate unwanted variation factors based on spike-in reads by running the command *RUVg(k=1)* <u>(Risso et al. 2014)</u>. The factors from the RUV method were applied in both DESeq2 and EdgeR analysis as one factor in the design matrix along with the treatment factors <u>(Risso et al. 2014)</u>.

The distribution of transcripts before and after normalization was visualized as an RLE plot using the *plotRLE()* function in the EDASeq package (Risso et al. 2011). The PCA plot was created by the *plotPCA()* function in the EDASeq package (Risso et al. 2011).

### 5. Differential expression analysis

DESeq2 and EdgeR were conducted on normalized read counts (Robinson, McCarthy, and Smyth 2010; Love, Huber, and Anders 2014) to obtain differentially expressed genes between times of day and between temperature conditions in R version 4.2.3. Genes identifiedas differentially expressed had an absolute log_2_ fold change higher than 0.5 and the FDR less than 0.05. Visualization including volcano plots, dot plots, Venn diagrams and heatmap plots was created by EnhancedVolcano (version 1.4.0) (Blighe et al. 2022), ggplot2 (version 3.4.2), VennDiagram (version 1.7.3) (Chen and Boutros 2011), and pheatmap (version 1.0.12) packages in R version 4.2.3.

#### 2.4.6 Functional analysis with MapMan and GO Enrichment analysis

MapMan analysis (version 3.5.1R2) was performed on the DEGs obtained from the Median of Ratio and spike-in, Nu*Delta, methods in DESeq2 (Thimm et al. 2004). The sorghum locus IDs that begin with ‘Sobic’ from the current annotation version (3.1.1) were converted to the locus IDs beginning with ‘Sb’ to be compatible with the sorghum database in MapMan. The ID conversion list was downloaded from the SorGSD database (Liu et al. 2021).

The R package ‘topGO’ (version 2.48.0) was used to determine enriched biological process GO terms <u>(Alexa and Rahnenfuhrer 2022)</u>. Sorghum annotated GO terms were from AgriGO v2.0 <u>(Tian et al. 2017)</u>. The algorithm ‘weight01’ with Fisher statistics was used to calculate the significant levels.

## Accession numbers

Sequencing data can be found at the National Center for Biotechnology Information Sequence, Read Archive (SRA) database (accession number TBD).

## Supplemental data

Supplemental Table 1 Read count in the synthetic RNA-Seq dataset for demonstrating the effect of global transcription change.

Supplemental Table 2 Read count in the synthetic RNA-Seq dataset for demonstrating the effect of proportional changes of the RNA pool with similar library size across samples.

Supplemental Table 3 Number of DEGs and confusion matrix tables under the effect of proportional changes of the RNA pool.

Supplemental Table 4 Number of reads after pre-processing (adapter trimming and rRNA removal), genome alignment, and gene assignment.

Supplemental Table 5 Genes encoding the subunits of RNA polymerases were differentially expressed between AM and PM in control and chilling stress conditions according to the Median of Ratio and RNA spike-in normalization methods.

## Acknowledgements

Author contributions

A.R.L., and C.J.D conceived the project and designed the experiments. A.V., I.M.S., and S.V.K.J. grew plants and collected plant tissues. K.L. prepared samples for RNA sequencing, analyzed sequencing data, developed the normalization pipelines, generated the synthetic data sets and graphs, and performed the qRT-PCR. All authors discussed the results and reviewed the manuscript.

## Supplemental Figure Legends

**Supplemental Figure 1 RNA spike-ins captured difference in transcript abundance due to altered global transcription and altered proportional shares of transcripts.** RLE plots of synthetic gene read counts before and after Median of Ratio and spike-in normalizations in DESeq2 under altered global transcription (A) and (B, C, and D) the drastic change in gene expression in condition B (B, C, and D) with consistent library size (A, B, and C) and varied library size across samples (D).

**Supplemental Figure 2 Details of sorghum 3’ RNA-Seq libraries and the relationship between gene and spike-in read counts.** (A) Read distribution on the gene body. The gene body coverage was calculated by the RSeQC package (version 2.6.6) <u>(Wang et al. 2012)</u>. 3’ RNA-Seq contributed to reads mapped to the 3’ end of the transcripts. (B) Correlation between total spike-in reads and total filtered gene reads. Pearson correlation coefficient and p-value are shown in the plot. (C) The proportion of spike-in reads to total reads. The boxplot shows the median of spike-in proportion in four treatments. One-way ANOVA indicated that there was no significant difference (p-value > 0.05) in the mean of the proportions between treatments.

**Supplemental Figure 3 RNA spike-in normalization approaches identified more evening-upregulated genes in both control and chilling stress conditions.** (A and B) The number of DEGs between morning and evening under control (A) and chilling temperatures (B) from different normalization approaches in DESeq2 and EdgeR (See Materials and Methods for details about each normalization). Traditional normalization methods in DESeq2 and EdgeR are Median of Ratio and TMM, respectively. RUV-based approches use spike-ins to normalize for library size, but do not account for changes in global transcription. Genes with FDR < 0.05 and the absolute log_2_ fold-change > 0.5 were identified as differentially expressed genes. Bold y-axis labels highlighted methods utilizing external RNA spike-ins as normalization factors, and and italic labels indicated analyses in DESeq2. (C) Volcano plot displayed the log_2_ fold-change and p-value of 1370 genes identified as DEGs up-regulated in the AM by the traditional, Median of Ratio method in control conditions. Their -log_10_P and log_2_ fold-change values are plotted for their normalization with the spike-in normalization to visualize why they were not detected with the spike-in approach. Dot plots represent the example of genes that were not differentially expressed by the Median of Ratio method due to failed log_2_ fold-change (blue), failed p-value (green), and failed log_2_ fold-change and p-value (black). (D) Volcano plot displayed the log_2_ fold-change and p-value of 6142 genes uniquely identified as evening up-regulated genes with the spike-in normalization under chilling stress condition. Their -log_10_P and log_2_ fold-change values after the Median of Ratio normalization are plotted to visualze why these were not identified by the Median of Ratio approach. Dot plots represent the example of genes that were not differentially expressed by the Median of Ratio method due to failed log_2_ fold-change and p-value. (E) Volcano plot displayed the log_2_ fold-change and p-value of 2012 genes uniquely identified as morning up-regulated genes with the spike-in normalization under chilling stress. Their -log_10_P and log_2_ fold-change values are plotted for the Median of Ratio normalization. Dot plots represent the example of genes that were not differentially expressed by the spike-in normalization due to failed log_2_ fold-change and p-value.

**Supplementary Figure 4 RNA spike-in normalization approaches identified more chilling down-regulated genes in the morning but more chilling up-regulated genes in the evening.** (A and B) The number of DEGs between control and chilling stress in the morning (A) and in the evening (B) from different normalization approaches in DESeq2 and EdgeR (See Materials and Methods for details about each normalization). Genes with FDR less than 0.05 and the absolute log_2_ fold-change higher than 0.5 were identified as DEGs. Bold y-axis labels highlighted methods utilizing external RNA spike-ins as normalization factors, and and italic labels indicated analyses in DESeq2. (C to F) Heatmaps represent lists of significant GO terms (adjusted p-value < 0.01) of AM chilling down-regulated genes (C), AM chilling up-regulated genes (D), PM chilling down-regulated genes (E), and PM chilling up-regulated genes (F) from the Median of Ratio and spike-in normalization methods. Numbers inside the boxes are the numbers of genes and colors indicate the log_2_ of adjusted p-values.

**Supplementary Figure 5 Details of external RNA spike-in controls and spike-in normalization methods used in the sorghum 3’ RNA-Seq dataset.** (A) Concentrations and average observed read counts of detected and undetected ERCC transcripts grouped by experimental conditions. A red vertical line indicated a 2 amol/μL of ERCC controls. (B) Workflow of differential expression analyses with different normalization approaches (See Materials and Methods for details). Figure was created with Biorender.com.

