## Supplementary figures and images for "A normalization method that controls for total RNA abundance affects the identification of differentially expressed genes, revealing bias toward morning-expressed responses"

### Supplemental Figure 1

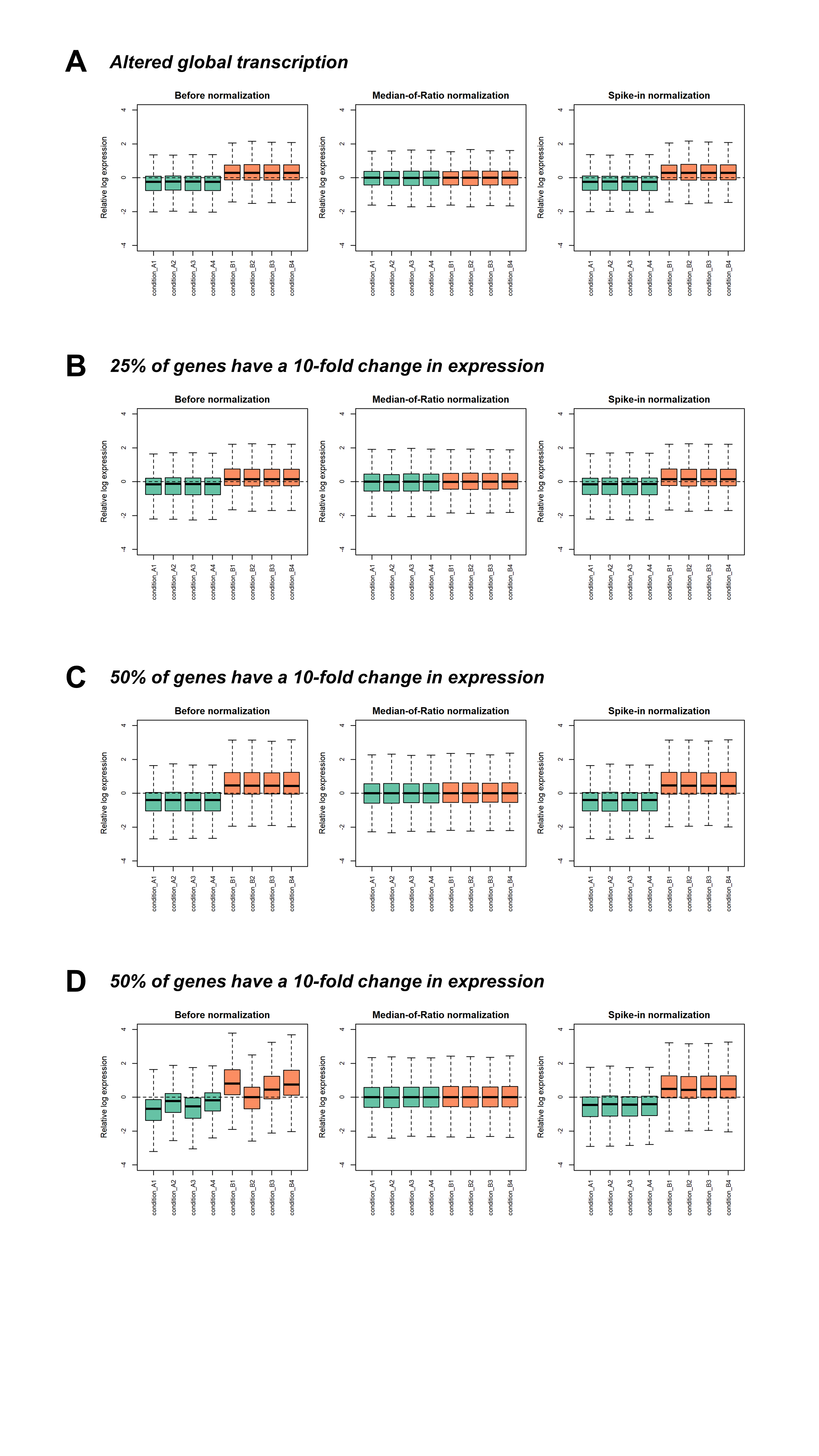

### Supplemental Figure 2

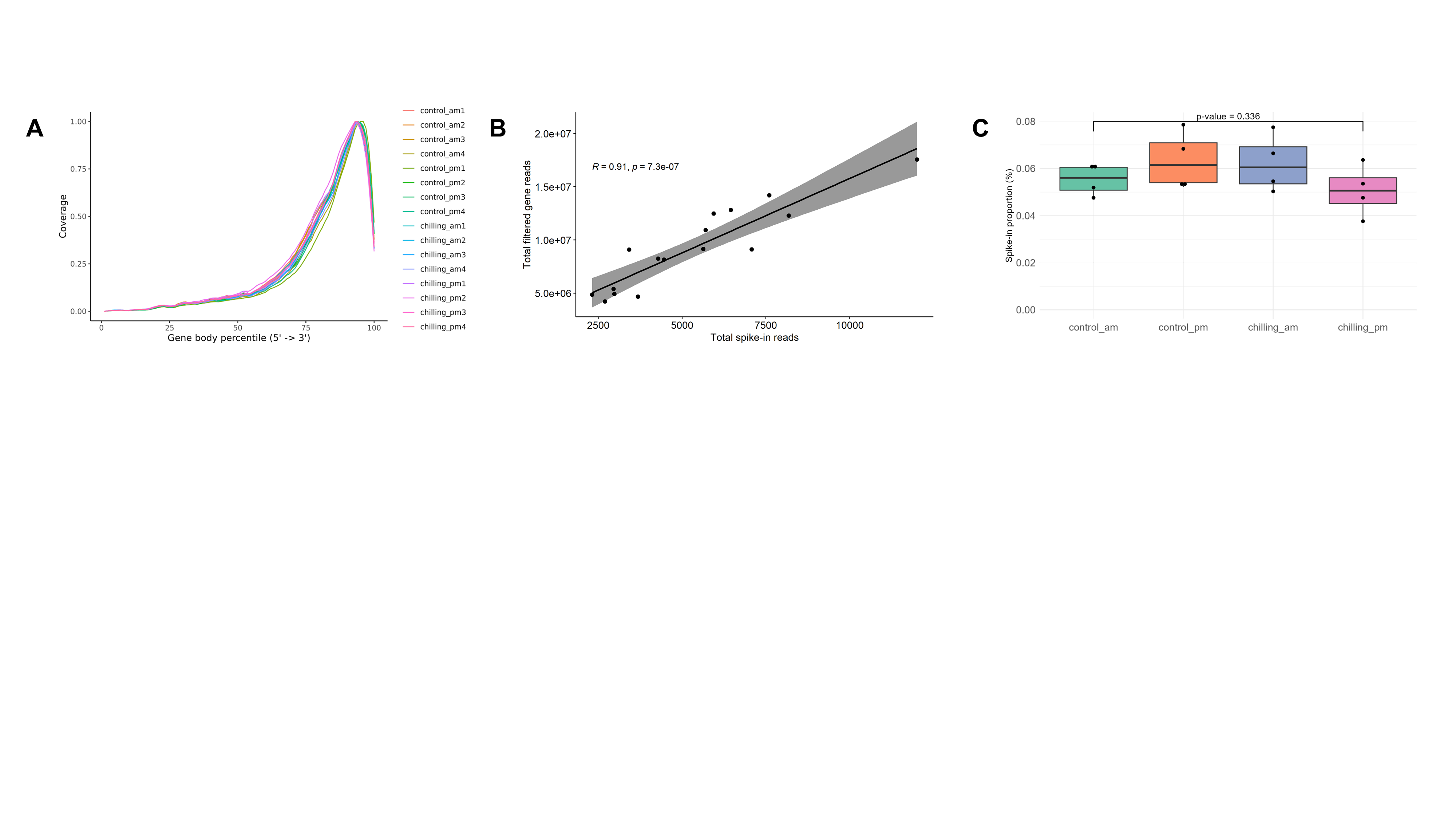

### Supplemental Figure 3

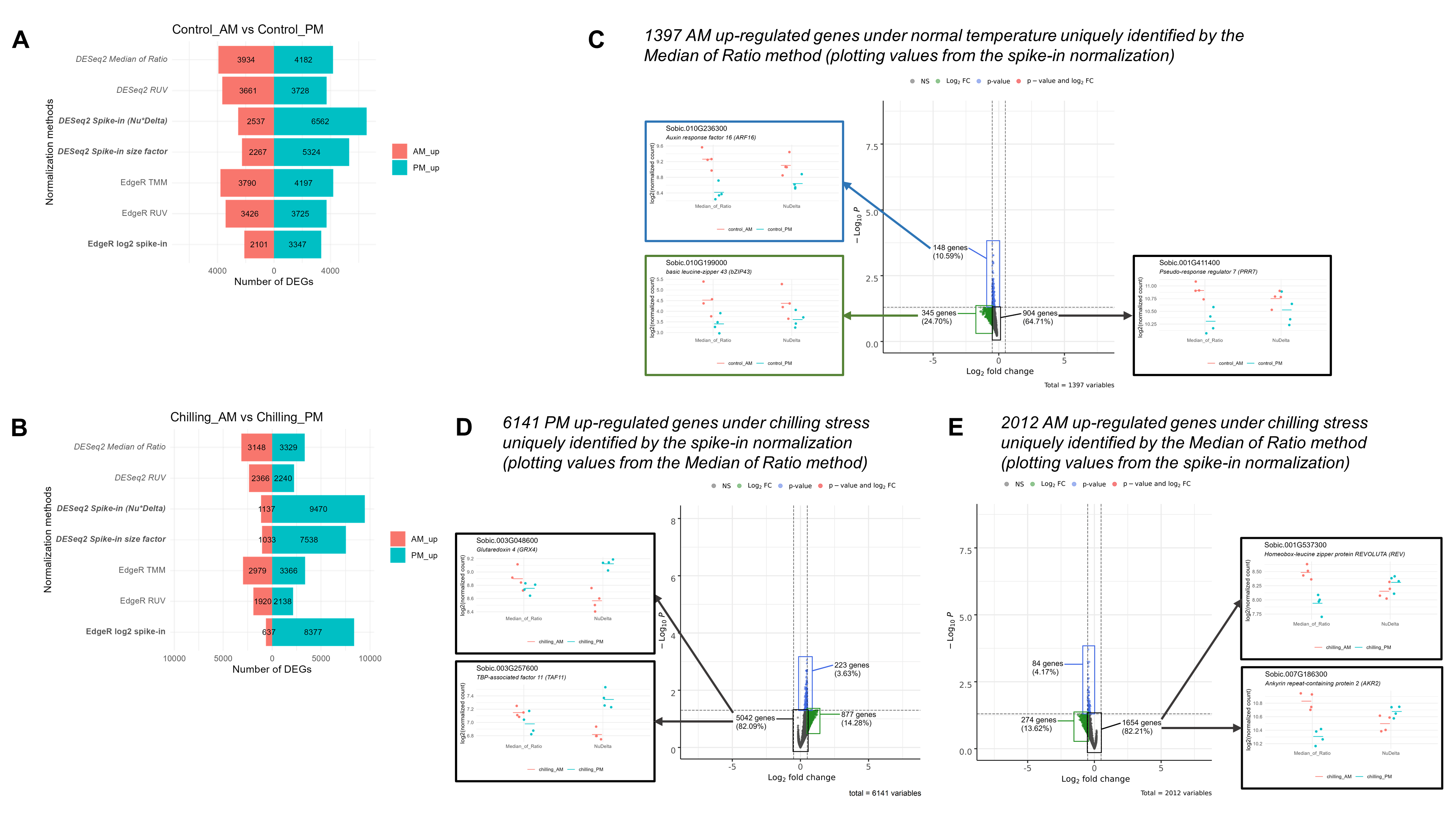

### Supplemental Figure 4

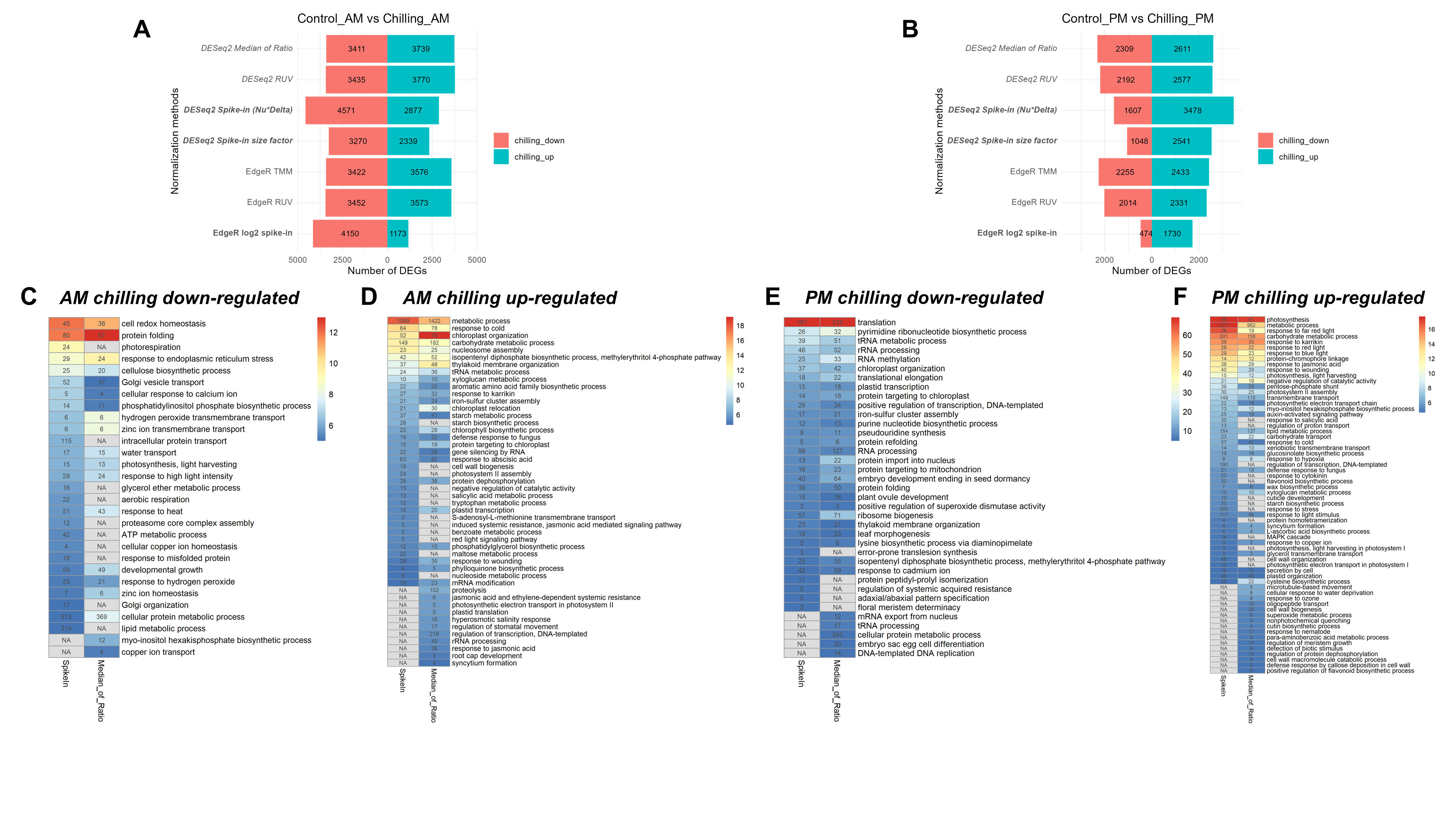

### Supplemental Figure 5

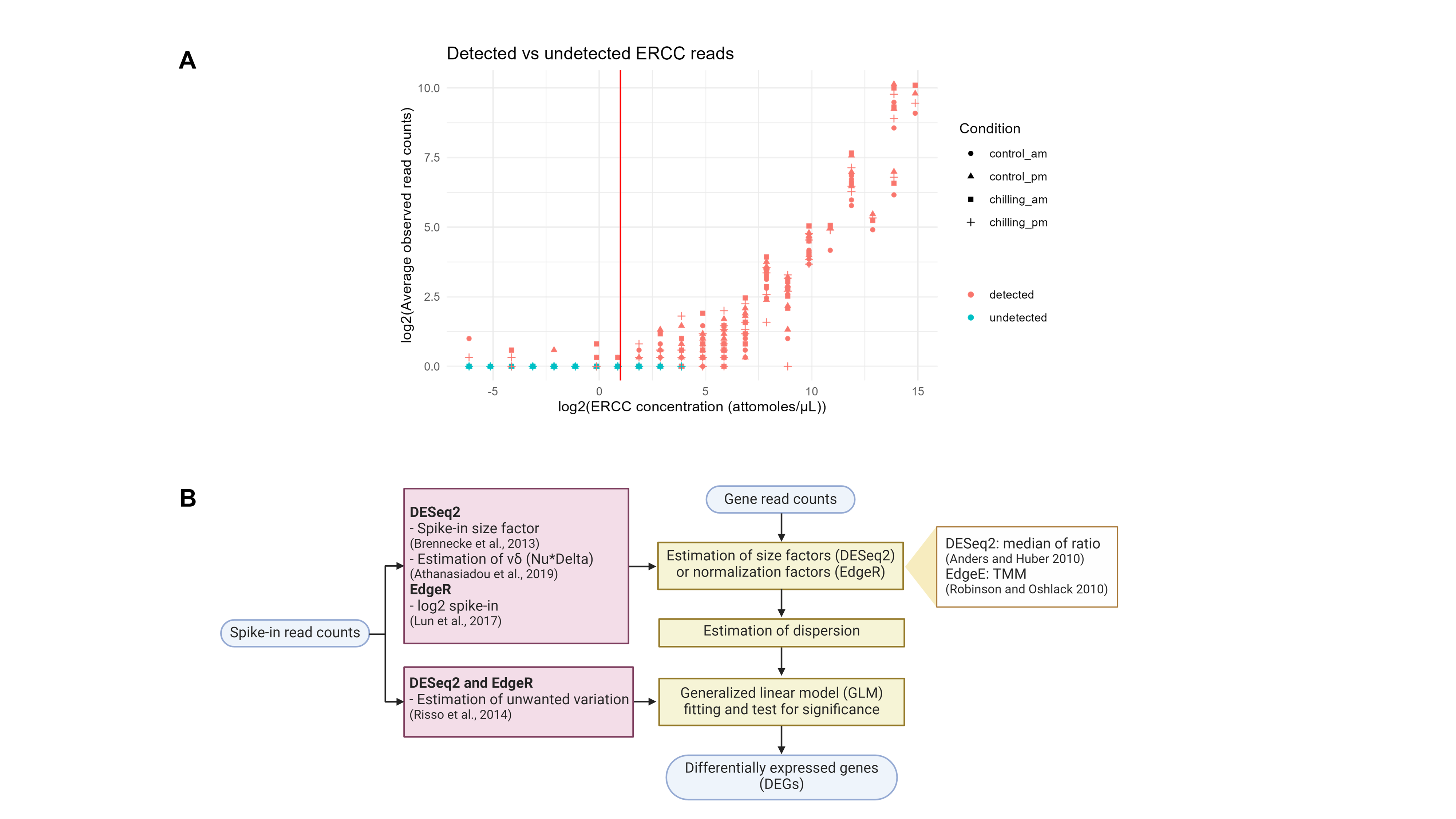
